# Plasmid Permissiveness of Wastewater Microbiomes can be Predicted from 16S rDNA sequences by Machine Learning

**DOI:** 10.1101/2022.07.09.499415

**Authors:** Danesh Moradigaravand, Liguan Li, Arnaud Dechesne, Joseph Nesme, Huda Ahmad, Manuel Banzhaf, Søren J Sørensen, Barth F Smets, Jan-Ulrich Kreft

## Abstract

Wastewater Treatment Plants (WWTPs) contain a diverse microbial community with high cell density. They constantly receive antimicrobial residues and resistant strains and, therefore, may offer conditions for the Horizontal Gene Transfer (HGT) of antimicrobial resistance determinants, transmitting clinically important genes between, e.g., enteric and environmental bacteria and *vice versa*. Despite the clinical importance, tools for predicting HGT are still under-developed. In this study, we examined to which extent microbial community composition, as inferred by partial 16S rRNA gene sequences, can predict plasmid permissiveness, i.e., the ability of cells to receive a plasmid through conjugation, for microbial communities in the water cycle, using data from standardized filter mating assays using fluorescent bio-reporter plasmids. We leveraged a range of machine learning models for predicting the permissiveness for each taxon in the community, translating to the range of hosts a plasmid is able to transfer to, for three broad host-range resistance plasmids (pKJK5, pB10, and RP4). Our results indicate that the predicted permissiveness from the best performing model (random forest) showed a moderate-to-strong average correlation of 0.45 for pB10 (95% CI: 0.42-0.52), 0.42 for pKJK5 (0.95% CI: 0.38-0.45) and 0.52 for RP4 (0.95% CI:0.45-0.55) with the experimental permissiveness in the unseen test dataset. Predictive phylogenetic signals occurred despite these being broad host-range plasmids. Our results provide a framework that contributes to assessing the risk of AMR pollution in wastewater systems. The predictive tool is available as a an application under https://github.com/DaneshMoradigaravand/PlasmidPerm.

## Introduction

Antimicrobial resistance (AMR) is a global threat causing an increasing burden across healthcare settings worldwide [1, 2]. Wastewater treatment plants (WWTPs) serve as key monitoring and control points connecting different community and hospital sewers with receiving aquatic environments [1, 3]. WWTPs therefore receive antibiotics from human consumption in the community and hospitals [4], which may not diminish even after the treatment process [5] and thus contribute to the residual antimicrobials in the environment [6]. Besides antimicrobials, and probably more importantly, the mixed sewage contains diverse antimicrobial resistant strains, which often carry their resistance genes on plasmids. These sites, therefore, serve as key points in the dissemination network of AMR determinants [7–9].

The evolution of AMR is driven by a combination of genetic mechanisms, i.e., mutations and Horizontal Gene Transfer (HGT). One of the major mechanisms of HGT is via conjugation of plasmids or integrative conjugative elements, which is thought to transfer ARGs between both closely and distantly related lineages in microbial communities such as those in WWTPs [8, 10]. In WWTPs, commensal and pathogenic strains of human origins are mixed with environmental bacteria, and the high cell density and ability to grow can provide conditions for genetic exchange of mobile genetic elements carrying antimicrobial resistance genes (ARGs), facilitated by subinhibitory concentrations of residual antimicrobials in WWTPs [11].

The 16S rRNA gene is an essential, conserved gene across all bacterial and archaeal lineages. However, variation in the hypervariable regions (V1–V9) of the gene allows differential identification of taxa, though only at the genus level if short-read sequencing is used [12, 13]. Despite these limitations, the feasibility and cost-effectiveness of 16S rRNA gene amplicon sequencing has promoted its popularity in microbiome studies. It has allowed detecting the association of taxonomic community composition with a range of ecological dynamics and habitat characteristics, e.g., disease association [14, 15] or ecological status [16].

Recent efforts have characterized the possible extent of genetic exchange in WWTPs by leveraging the strength of 16S rDNA amplicon sequencing and *in vitro* mating assays where fluorescent tagging enables collecting transconjugants by fluorescence-activated cell sorting (FACS) followed by sequencing [17–20]. Using this approach, a recent in-depth analysis of activated sludge microbial communities to serve as recipients of HGT of three broad-host range multidrug resistance plasmids was carried out [21]. The ability of a recipient cell to receive and maintain a plasmid (at least for a short duration) is referred to as its permissiveness [21]. Using the reporter system enables direct quantification of plasmid permissiveness for all recipients in a community and thus define the host range of the plasmid in that community. This study did not detect a phylogenetic signal in permissiveness, leading to the conclusion that translating permissiveness of a bacterial group to other phylogenetically similar groups in the WWTP community would not be valid [21]. Another study found permissiveness to vary strongly across the recipients’ taxa [22]. These studies, however, did not examine the predictive power of sequence markers for plasmid permissiveness.

In the context of microbial molecular ecology studies, machine learning approaches have proven an effective tool for predicting phenotypic features from genomic biomarkers such as antimicrobial resistance, host of isolation, bacterial growth and virulence [23, 24]. The models can predict the features without any prior knowledge about the mechanisms, by learning complex, nonlinear and high-order phylogenetic signals in a training dataset of labeled sequences to be used for rapid detection of the trait in unseen data [25, 26]. Among the various models that were employed in these studies, ensemble methods, e.g. random forests and gradient-boosted trees, were consistently found superior to other methods including linear models. These models combine multiple weak learners to account for overfitting while utilizing high order interactions between predictive features for prediction. While several phenotypic features are used as labels in the models, machine-learning algorithms have not been employed for the prediction of HGT features.

In this study, we leveraged machine learning approaches to assess the degree to which taxonomy (amplicon sequencing of the V3-4 hypervariable regions of the 16S rDNA) can predict permissiveness of recipient communities for broad host-range plasmids from *in vitro* conjugation mating assays. Predictive power for narrow host-range plasmids should be very high but was not tested as data were not available. We employed a range of machine learning regression methods for an alignment free (i.e., kmer representation) input of sequence data. Our results indicate that the sequence data predicted the permissiveness for three broad host-range AMR plasmids with an average accuracy of 0.63 for the correlation of predicted and actual values on the unseen data. We identified a set of predictive kmer sequences and how these are distributed across diverse host taxa. These results suggest that permissiveness can be partially predicted without full genome sequencing, and can be exploited for predicting the ability of resistance plasmids to spread into particular recipients or communities of recipients in environments. Since there are far too many environments, species and plasmids for field-scale surveillance of HGT, focusing experimental effort on generating data for machine learning to predict permissiveness is proposed here as a solution.

## Methods

### Study design and sampling

To have a comprehensive microbiome collection from different time points and locations in the water cycle, we retrieved samples from one WWTP in the UK and one in Denmark in 2017 and 2018. Samples from the UK and Denmark were taken at different locations along the sewage treatment process (Figure S1). This included residential and hospital sewers, the point where they mix, the WWTP influent, after the primary and secondary settlers, in the biological treatment stage, after the tertiary filter and upstream and downstream of the effluent entering the receiving river. The sampling dates and locations are provided in Supplemental Table S1.

### Experimental filter mating assay

We employed the Solid Surface Filter Mating Assay [20, 27] to measure the permissiveness of water cycle microbial communities toward three typical conjugative plasmids. Cell suspensions of donor strain and WWTP recipient community were mixed at 1:1 cell ratio and immediately filtered. The filter was then placed on an agar-solidified synthetic wastewater medium. After incubation (48 h at 25°C) and GFP maturation (48 h at 4°C), transfer events were detected by epifluorescence microscopy and transfer frequency was quantified as the ratio of conjugation events (CE) detected as GFP microcolonies to the original number of recipients (R) in the sample (CE/R). Here, recipient cells were untagged bacteria from the environmental sample. We used *E. coli* MG1655 as a donor (chromosomally tagged with mCherry expressed from the constitutive promoter pLpp) carrying one of the three plasmids, pKJK5 (IncP-1ε), pB10 (IncP-1β), and RP4 (IncP-1α) [28] (plasmids were tagged with pLac-gfp repressed by a chromosomal lacI^q^). Thus, donors had red fluorescence, recipients none and transconjugants green fluorescence.

### Sorting and sequencing

For each filter mating, cells were recovered from the filters and transconjugants and recipients were gated and sorted by fluorescence-activated cell sorting if they were of bacterial size (using the forward scatter detector) and presented green fluorescence (transconjugant pool) or an absence of red fluorescence (recipient pool), respectively, as described previously [18, 29]. Sorted cells were subjected to DNA extraction. The hypervariable regions V3–V4 of the 16S rRNA gene were amplified using primer set 341F/806R and went through paired-end sequencing on an Illumina MiSeq platform. We analyzed the paired-end reads of 16S rRNA gene amplicon sequencing using the DADA2 pipeline to obtain amplicon sequence variants (ASVs) [30, 31]. We excluded sequences that were longer than 430 bp. his resulted in 2272 unique ASVs.

### Calculating permissiveness

We define permissiveness as the ability of a host cell (identified as an ASV) to receive and maintain a plasmid (at least for a short duration). As estimating ASV-specific permissiveness is complicated by the (potential) growth of both transconjugants and recipients during mating incubation, we calculated apparent permissiveness (AP). It is defined as the ratio of the relative abundance of an ASV in the transconjugant pool to the corresponding recipient pool [29]. AP thus accounts for the fact that the abundance of an ASV in the transconjugant pool partly depends on its abundance in the recipient pool. When calculating permissiveness, we assigned a count of one to an ASV in the recipient pool if it was absent while it was present in the transconjugant pool because at least one recipient must have been there if a transconjugant was detected, so the ASV must have been missed, e.g., because the sampling was not sufficiently exhaustive. Permissiveness values reported throughout the paper are apparent permissiveness values. The sequences and permissiveness values are detailed in Supplemental Table S1.

The replicates for the pKJK5 plasmid permissiveness measurements showed a correlation of 0.74 (Spearman’s rank correlation coefficient) between the values for the same taxa, which was used as a proxy for the maximum possible correlation value. To account for the measurement error, we divided the measured correlations of permissiveness values by this proxy.

### Machine Learning platform

We developed a machine learning platform for predicting permissiveness and identifying predictive sequence signals. We opted for two approaches: point prediction for exact point prediction and interval prediction to account for the uncertainty. The point prediction platform invoked a baseline model of lasso regression and two ensemble models of gradient boosted regressors and random forest regressors. We used the built-in functions in the sklearn 1.0.2 library for that purpose [32]. We scaled the response variable prior to feeding into the machine learning algorithm. We opted for three-fold cross validation and split the data into 80% and 20% training/validation and test datasets. The machine learning models were tuned by taking a grid search approach. For the lasso model, we tuned the L2 regularization penalty term, by assessing the values (0.0001, 0.001, 0.01 and 0.1). For gradient boosted regressors, we tuned the key parameters of tree depth parameter for values 1, 3 and 5, and the number of iterations of 50, 100 and 200. For random forest, we tuned the key tree-related parameters of the number of trees (20, 50 and 100) and the tree depth (1, 3 and 5). We identified the best performing model as the model with the smallest average minimum absolute error (MAE). To obtain error intervals for the prediction, we repeated the training/test model for ten random train/test data splits. To assess the performance of the tuned model, we computed Spearman’s rank correlation coefficient or Spearman’s ρ between the predicted and actual data.

Besides point prediction of the output, we used random forest models to obtain prediction intervals. The intervals were computed based on the predictions from all the learners (trees) in the final tuned ensemble model (random forests). The models account for the uncertainty both in fitting the model and in the sampling and sequencing. In assessing the model performance, we considered the true detection rate or coverage, which corresponds to the proportion of test observations that were covered by the prediction intervals at different confidence levels. We used the rand_forest() in R and compared the intervals with the dispersion around the mean (mean absolute difference) and measurement error range.

We employed an alignment free approach for generating predictor features [33]. The approach better accounts for highly divergent sequences and allows the application of the trained model to unseen data without the need for time-consuming multiple alignment and training steps. To this end, we enumerated the kmers of increasing sizes of 5 to 8 (a larger kmer size was not found to improve the prediction performance). This resulted in a matrix with numbers indicating the frequency of the kmers in each sequence. We scaled the values using a min-max scaler prior to feeding them to the machine learning models. If a transconjugant’s sequence was found in more than one site, we averaged the measured apparent permissiveness values for that sequence across the sites. In random forest models, we measured the importance of features, i.e., describing which features are relevant, as the decrease in node impurity weighted by the probability of reaching that node. The node probability can be calculated as the number of samples that reach the node, divided by the total number of samples. The higher the value, the more important the feature. To robustly identify important predictive features, we repeated the prediction on ten random splits of train/test data and extracted the important features. We aggregated the results across replicates and kept the features that were found important across all ten replicates in the training dataset. We then filtered the top five percent of the features according to their average ranks. We excluded kmers if they were found in longer predictive kmers. We deployed the trained and tuned models as web and command line applications, which estimates the permissiveness for any user input 16S rDNA sequence. The tool is available in https://github.com/DaneshMoradigaravand/PlasmidPerm.

### Taxonomic classification analysis and association analysis

We examined the predictive information contained in the taxonomic clusters inferred from 16S rDNA ASVs of transconjugants for permissiveness, to understand whether elevated/lower permissiveness is linked with particular clades (lineages) or whether it is a trait emerging across multiple taxonomic groups in the transconjugants. Therefore, we employed BAPS, Bayesian Analysis of Population Structure [34], as implemented in the R package rhierbaps [35], to analyze transconjugant communities. Although BAPS is used to identify population sub-structures for one species, we adopted it for finding taxonomic clusters in 16S rRNA data. Here the clustering ASV features correspond to panmictic BAPS clusters. We screened increasing the number of iterations to capture taxonomic classifications at different levels. After identifying the groups, the associations to the taxonomic groups were then converted into vectors using One Hot encoding before feeding into the machine learning framework. We trained and tuned the model using the same platform as above and assessed the performance of the model on the held-out test dataset. For identifying important features, we repeated the prediction on 10 random splits of training/test datasets and kept the features that appeared in all replicates.

To identify the kmers that were not associated with a particular clade, we enumerated all possible kmers with increasing sizes of 5 to 12 kmers and created an input matrix with zero and one values, indicative of the absence and presence of the kmers, respectively. We then binarized the response variable (plasmid permissiveness) according to the median value. We used the software Scoary [36] for examining the association of kmers and their respective plasmid permissiveness values. Only kmers with Bonferroni corrected p-values smaller than 0.05 and p-values corrected for population structure, i.e., best and the worst possible *p* values, smaller than 0.05 were kept.

### Evaluation of prediction model with simulated data

We simulated sequences to understand the impact of mutation rate, sample size and strength of correlation between predictive kmer and permissiveness on the prediction accuracy and extent of overfitting. We employed Simbac [37] and simulated collections of 150bp kmers with 100, 200, 500 and 1000 isolates and under mutation rates of 0.01, 0.5, 1 and 10 mutation per time unit. We introduced the predictive kmer by randomly selecting a kmer with frequency >0.1 for 20 times. We attributed permissiveness to isolates with and without the kmer by introducing a coefficient value (*λ*), which determined the sampling from the baseline pKJK5 plasmid permissiveness. For isolates with the kmer, we drew a random value from the distribution of permissiveness values for the pKJK5 plasmid truncated between the value of *μ*/*λ* and the maximum permissiveness value, where *μ* is the mean of the permissiveness distribution for pKJK5. For isolates lacking the concerned plasmid, we drew permissiveness values from the distribution of permissiveness values for the pKJK5 plasmid truncated between the selection coefficient value of *μ* /*λ* and the minimum permissiveness value. Thus, by increasing the absolute values for 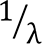, we increased the correlation between the predictive kmer and permissiveness and we therefore denote the correlation coefficient as 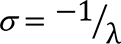. We then fed the simulated data into the predictive random forest pipeline, as detailed above, and computed the Spearman correlation between actual and predicted values for the held-out dataset.

## Results

We aimed to use the information in the 16S ASVs to predict plasmid permissiveness, i.e. the ability of cells to receive a plasmid through conjugation. The permissiveness values were obtained from filter mating assays for samples retrieved in 2017 and 2018 from one WWTP with an activated sludge process in Odense, Denmark, and one WWTP with trickling filters in Durham, UK (detailed in [38]). These were receiving residential and hospital sewage. Sampling locations are given in Figure S1A.

Overall, plasmid permissiveness showed a moderate positive correlation across transconjugant ASVs from different WWTPs, sampling sites and dates with a mean Spearman’s correlation of 0.49 (range:0.11, 0.73), which after correcting for the measurement error (see Methods) increased to 0.68 (Figure 1B). Except for 13 out of 180 pairwise comparisons of conditions, the correlation was significant (p-value<0.01, Spearman’s correlation test) (Figure 1B). The results suggest that the reproducibility of sampling the communities and the experimental procedure was somewhat limited at different time points and sites: For samples taken from the same site at the same time (replicates for sites 1A and 1B in Figure 1A) and for samples taken from the same sites at different time points in Denmark, average correlations of 0.68 and 0.75 were found for pKJK5 permissiveness values, respectively (Figure 1A).

**Figure 1.**
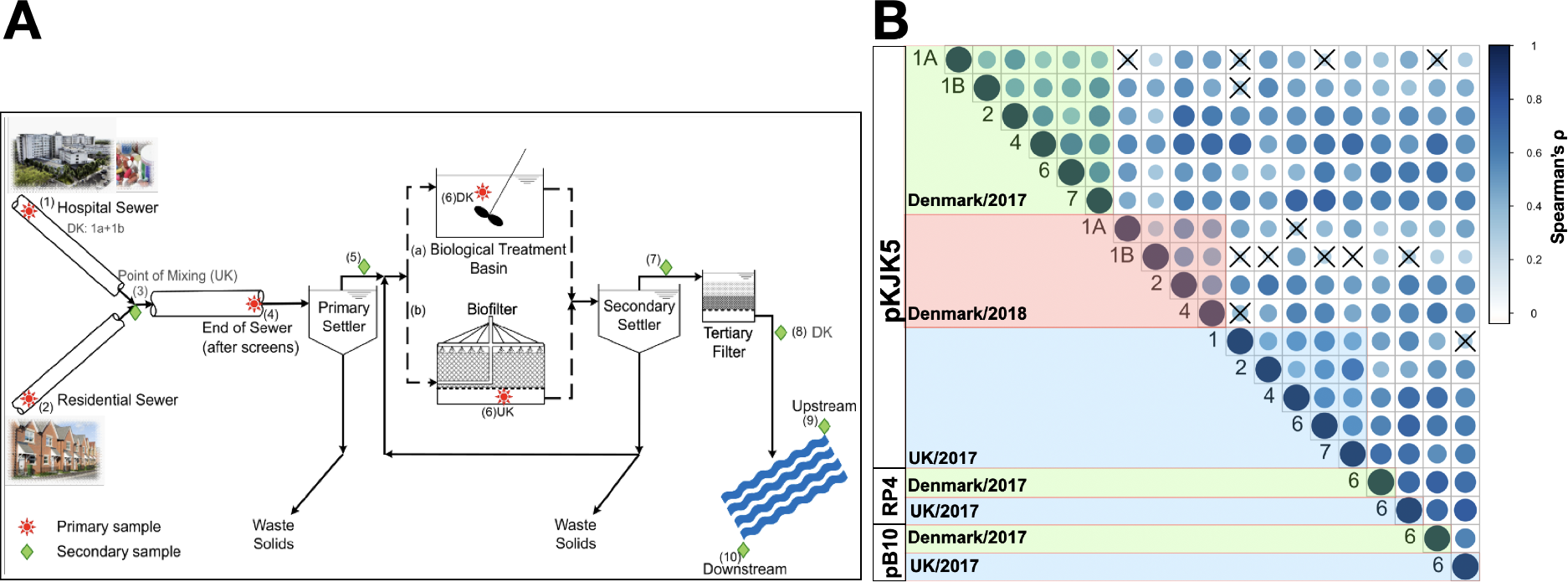
**(A)** The sampling points across the urban water cycles, note differences between the WWTPs in the UK and Denmark. **(B)** Correlations between plasmid permissiveness values at different sites, dates and plasmids are all positive. The cross signs label insignificant correlation (p-value < 0.01 from Spearman’s correlation test).

The measurements from the UK and Denmark form distinct clusters (Figure S1A), this may reflect different sewage composition, environmental conditions or treatment processes. Despite these differences between countries, the correlation between permissiveness across different plasmids was high and no clustering of measurements according to the plasmid type was apparent (average Spearman’s correlation of 0.68 across measurements for different plasmids) (Figure S1B and S1C), pointing to common mechanisms underlying permissiveness in recipient cells. To minimize the impact of time and location of sampling on the performance of the predictive models, we aggregated all permissiveness values for each plasmid.

We fed the permissiveness for the three plasmids as dependent training data into the predictive models to make point predictions of permissiveness. The models comprised a baseline regularized lasso regression model, a random forest and a gradient boosted regressor, which were trained on predictor features, i.e., the counts of kmers with different sizes (Figure 2). The results indicate that the ensemble models, i.e., random forest and gradient boosted regressor models, outperformed the lasso model in 10 out of 12 prediction settings, suggesting that accounting for nonlinear interactions improved prediction. Between the ensemble models, the random forest model was superior with the best average accuracy of 0.45 for pB10 (95% CI: 0.42, 0.52), 0.42 for pKJK5 (0.95% CI: 0.38,0.45) and 0.52 for RP4 (0.95% CI:0.45,0.55) across kmer values (Figure 1). After accounting for the measurement correlation of 0.74 (see Methods), the average accuracy can increase to 0.63 for pB10, 0.58 for pKJK5 and 0.72 for RP4. The extent of overfitting, i.e., the difference between the accuracy for the training and test datasets, did not vary for different models and kmers (Figure 2). To understand the impact of input transconjugant numbers on mitigating the overfitting, we repeated the prediction on down-sampled training datasets (Figure S2). We found that the increase in the numbers of transconjugants steadily improved the accuracy in the test dataset and reduced the overfitting up to the training size of 50% of the full size, beyond this, the improvement in accuracy leveled off (Figure S2). This finding suggests a significantly larger training dataset is required to further improve the prediction accuracy.

**Figure 2.**
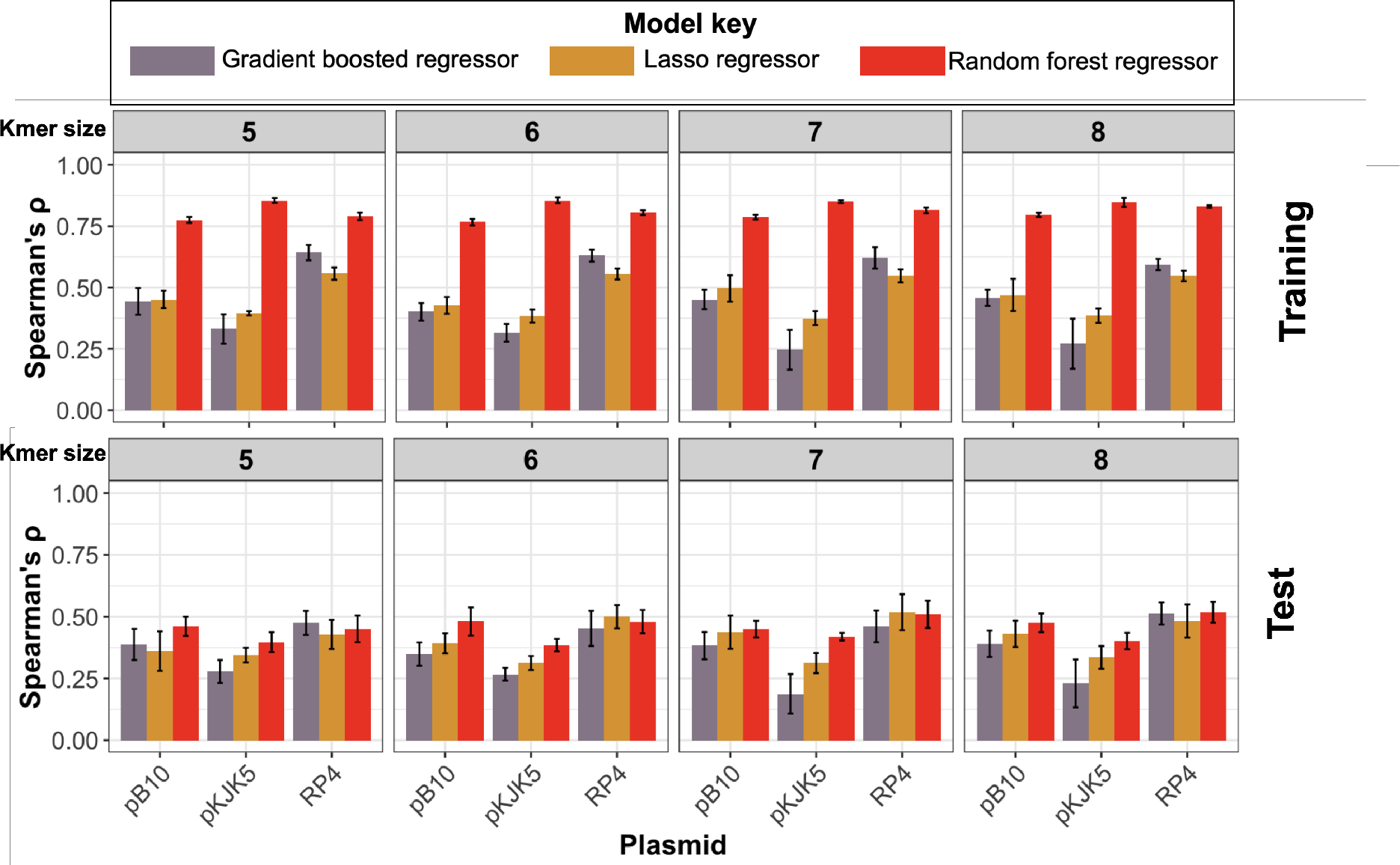
The accuracy of trained models for prediction of plasmid permissiveness for three plasmids (pB10, pKJK5 and RP4) and for gradient boosted, lasso and random forest regressors, in the training and test dataset. The error bars show 95% confidence intervals for models trained on ten random training/test splits.

As pointed out above, the differences in permissiveness values for two replicates indicated an uncertainty in the sampling and measurement, which led us to examine the random forest models that can yield a prediction interval. We therefore computed the associated uncertainty for the random forest model in the form of prediction intervals and then compared these prediction intervals with the measurement error and dispersion. Figure 3A shows the intervals containing 99% of the predictions around the point predictions. For the pKJK5 plasmid and a 99% interval, which is 0.57× and 6× the dispersion around the MAD (mean absolute difference) and measurement error range, respectively, the trained model correctly yielded a detection rate of 0.96 for the training and 0.94 for the test dataset (Figure 3A,B). For the pB10 (RP4) plasmids, the detection rate (coverage) of the predicted interval was lower and stayed at 0.64 (0.78) for the test data set at intervals that equaled 2.07× (3.5×) the mean absolute difference. As expected, with narrower prediction intervals, the coverage (detection) rate of the interval steadily decreased, however, the extent of overfitting remained low (Figure 3B). For pKJK5, with a prediction interval equals the average measurement errors, a detection rate of 0.67 on the test dataset was observed. For prediction intervals that equaled the dispersion of the data around the mean, the intervals included 57% of the permissiveness values for pKJK5 (Figure 3B). These values were 68% and 55% for predicting pB10 and RP4 permissiveness, respectively. Altogether, these results demonstrate the strength of the random forest model in capturing the uncertainty in measurements, which extends its applicability. However, resolving the uncertainty introduced by the model remains an open challenge.

**Figure 3.**
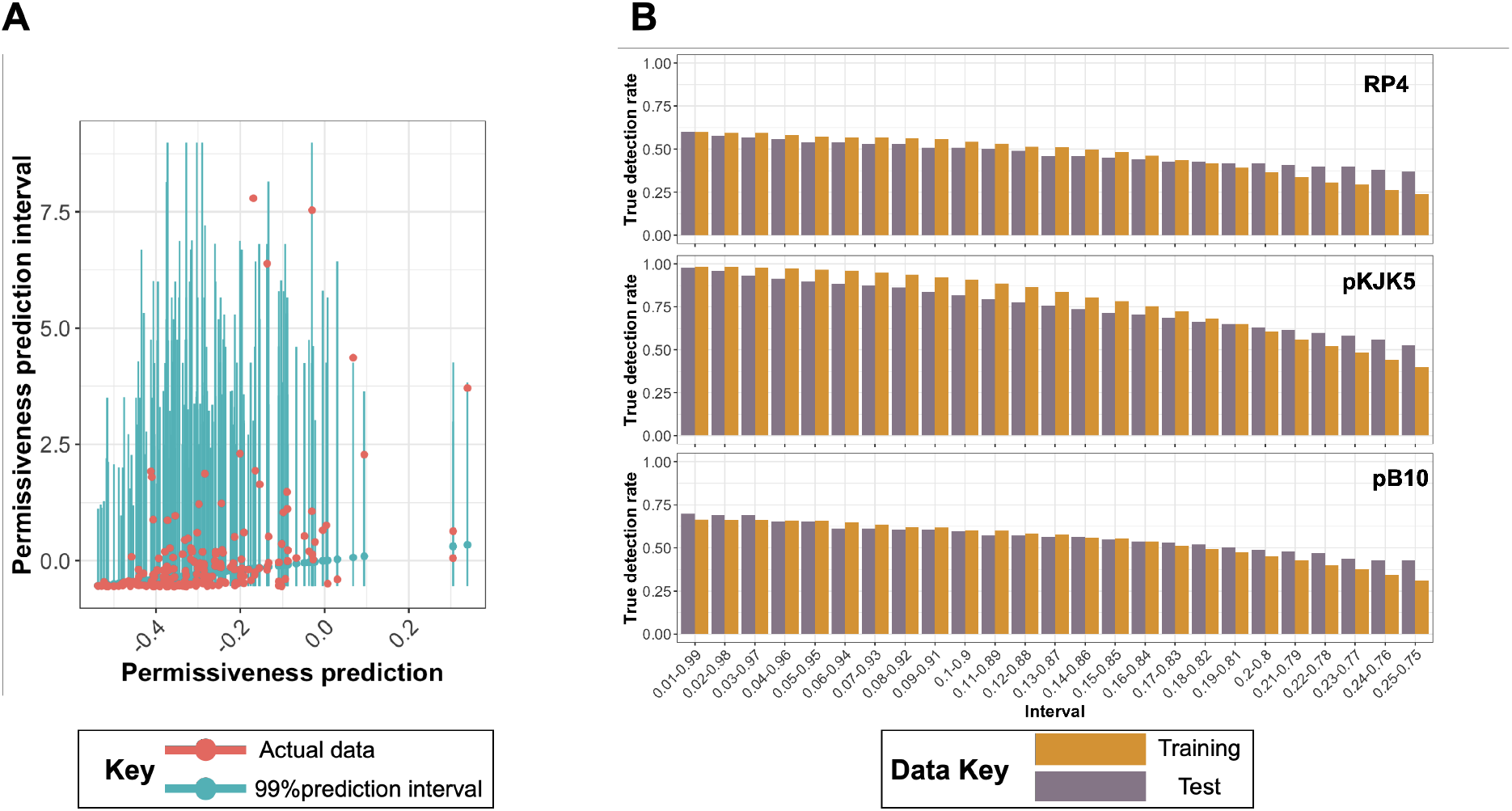
**(A)** Interval prediction for plasmid permissiveness with the random forest algorithm. Intervals contain 99% of predictions for permissiveness for the pKJK5 plasmid. **(B)** The true detection rate for different interval values for the pKJK5, RP4 and pB10 plasmids in the training and test datasets. The intervals on the X-axis contain the portions of data that fall between the upper and lower limits, e.g., the 0.01-0.99 range includes predictions greater than 1% and smaller than 99% of the values predicted by the model. True detection rate was defined as the relative frequency of data points that fell in the interval. The interval value of 99%, corresponds to 0.57×, 2.07× and 3.5× the dispersion around the mean absolute difference for the permissiveness distributions for pKJK5, pB10 and RP4 plasmids, respectively.

Although conjugation and maintenance of different IncP-1 plasmids is governed by shared mechanisms, genetic divergence in their transfer and regulatory regions has evolved [28], which may affect their transfer to and interaction with recipient cells, leading to differences in permissiveness. We therefore investigated the generalizability of models trained on one plasmid for predicting the permissiveness for a different plasmid. Prediction accuracy deteriorated for the test data for a different plasmid, as compared with prediction for the same plasmid, with an average correlation difference (Spearman’s ρ for training and test data for the same plasmid -Spearman’s ρ for training and test data for the different plasmids) of 0.32 for pB10, 0.14 for pKJK5 and 0.32 for RP4 plasmids (Figure S3). The partial drop in correlation when predicting permissiveness for other plasmids suggests plasmid specific differences in interactions with recipient cells.

Like plasmid types, our results indicated that the ‘country’ used for training data affects the accuracy of the prediction but it has to be emphasized that the WWTPs in the UK and Denmark use different biological treatment processes so the ‘country’ effect could be partially an effect of treatment processes. Models trained on UK transconjugant 16S rDNA sequences showed an average decreased accuracy (Spearman’s ρ for training data for both the UK and Denmark samples and test data for Denmark − Spearman’s ρ for training data for the UK and test data for Denmark) of 0.18, 0.15 and 0.19 for plasmids pB10, pKJK5 and RP4, respectively, when compared with models trained on mixed data (Figure S3).

In the same way, when we trained the model on all but one site and assessed the performance on the excluded site, the accuracy was lower than the accuracy when mixed data was used (average decrease in accuracy of 0.13 for Spearman’s ρ for training and test data for all sites mixed − Spearman’s ρ for training data for all sites except for the excluded site and test data for the excluded site) (Figure S3C). Overall, these findings demonstrate the need for representative training datasets to eliminate confounding factors that degrade model performance.

We then investigated whether using taxonomic clusters can improve predictions of permissiveness. The accuracy of predictions based on clustering ASV features (panmictic BAPS clusters) turned out to be consistently lower than predictions based on kmers (Figure 3A). For different types of models, the models trained on BAPS cluster information had an average diminished accuracy of 0.06 for pB10, 0.23 for RP4 and 0.21 for pKJK5, compared with the best performing kmer-based prediction (Figure 4A). Nevertheless, we identified eight predictive BAPS clusters for pKJK5, which were positively linked with permissiveness values (p-value from Wilcoxon test < 0.01) (Figure 4B). The phylogenetic distribution of the BAPS clusters showed that these clusters occur across a wide range of taxa (Figure 4C). We compared the frequencies of taxa contained in predictive BAPS clusters with the baseline frequency of the same taxa to identify the enriched orders (Figure 5A). Previous studies reported a broad host range for the pKJK5 plasmid [18, 39, 40]. In line with these reports, the predictive BAPS groups fell under both Gram-negative and Gram-positive orders. Most notably, Sphingomonadales and Bacillales turned out to be overrepresented in the predictive groups. This finding highlights the importance of HGT in the evolution of these strains, as shown for phage-mediated gene transfer [41]. The Gram-positive significant orders included a wide range of orders, e.g., Clostridiales, Bacillales, Micrococcales and Corynebacteriales (Figure 5D). The enriched orders for pB10 and RP4 turned out to be like pKJK5 but with some unique orders. While we observed a sharing of predominantly orders for pKJK5 with pB10 and RP4 plasmids, e.g., Micrococcales and Sphingomonadales, some recipients orders, e.g., Proteobacteria, Xanthomonadales, Alteromonadales and Pseudomonadales (Figure 5D), appeared exclusive to pB10 and RP4 plasmids. These orders contain many pathogens, in which HGT plays a major role in driving the evolution of adaptation, AMR and pathogenicity in humans and plants [42, 43]. Altogether, these findings suggest the existence of plasmid-specific interactions and shared recipient features in certain taxonomic clusters.

**Figure 4.**
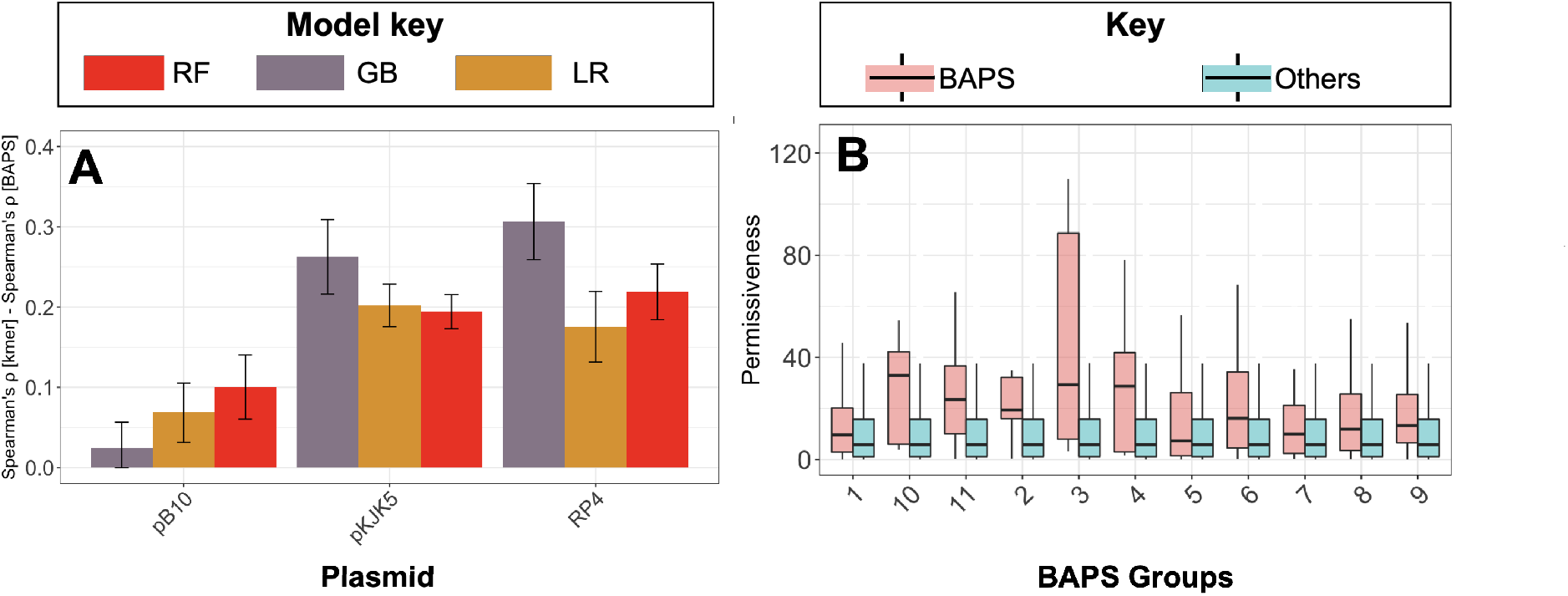
BAPS genetic cluster-based prediction. **(A)** The bars show the differences between the accuracies of the best performing models for the kmer-and BAPS-based models for different model classes. The terms “RF”, “GB” and “LR” stand for random forest, gradient boosting and lasso regressors, respectively. The error bars show 95% confidence intervals across 10 prediction runs with random test/training split. **(B)** The permissiveness values for ASVs belonging (red) or not-belonging (blue) to predictive BAPS clusters for plasmid pKJK5.

**Figure 5.**
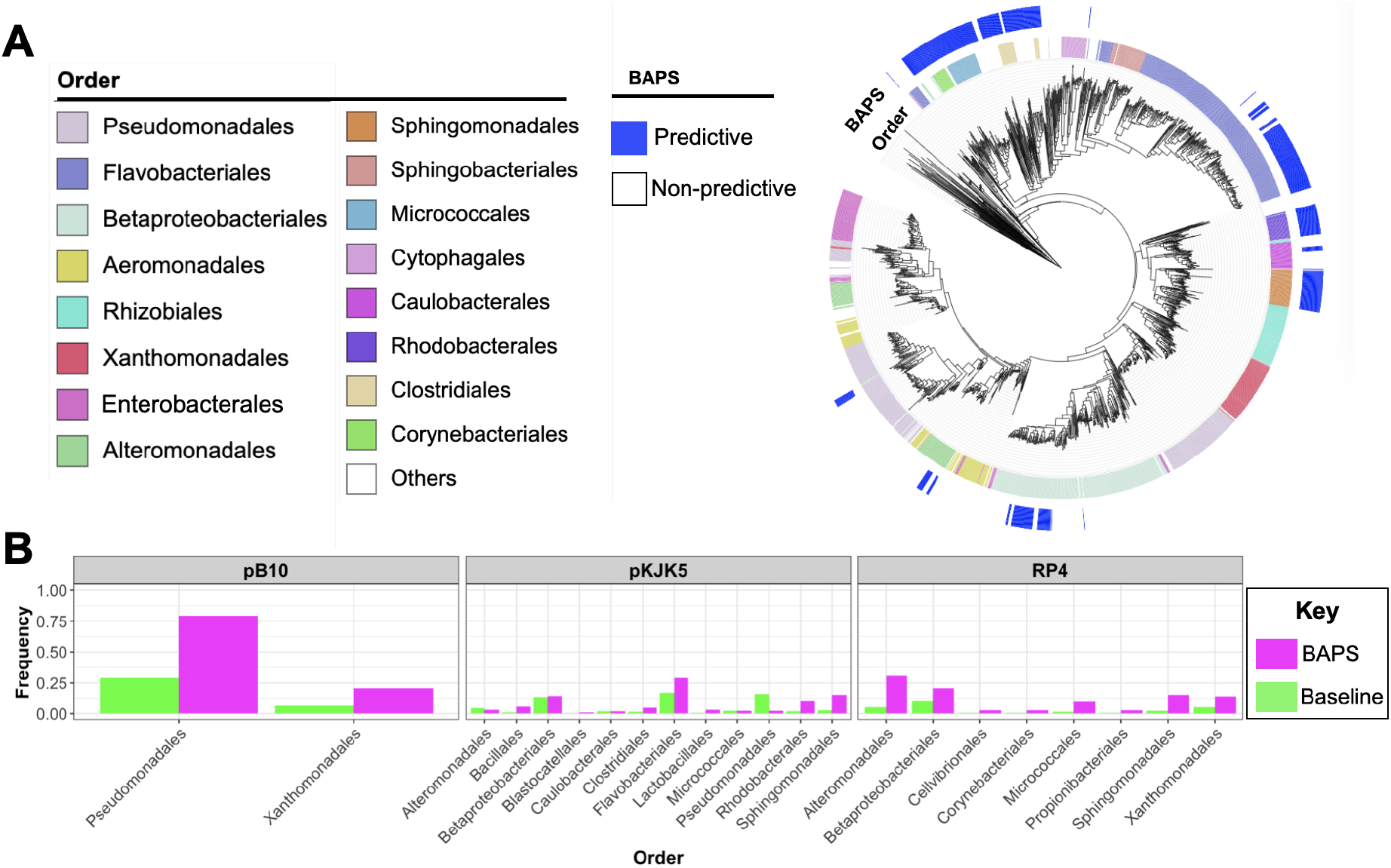
**(A)** The taxa distribution of predictive BAPS clusters for pKJK5. **(B)** The frequency of enriched orders in the predictive BAPS clusters for pKJK5, RP4 and pB10 plasmids. The green bars show the baseline frequencies in the recipient pool.

To examine the prediction information in kmers, we next identified kmers most predictive for permissiveness. The feature importance analysis for significant kmers pinpointed 83, 19 and 14 kmers, of which 79, 12 and 8 were positively linked with permissiveness for pKJK5, pB10 and RP4, respectively (Figures 6, S4). Like the BAPS clusters, we observed a distinctive distribution of significant kmers for each plasmid. However, we observed a higher discriminatory power between ASVs with and without the predictive kmers, as compared with ASVs from the predictive and non-predictive BAPS clusters for the plasmids (Figure 7). This implies that information in the predictive kmers accounted for a greater variance in the data, as compared to information in predictive BAPS clusters, pointing to multiple taxonomic signals within the ASVs that only kmer-based predictions used. This result is congruent with the better performance of kmer based predictions. The predictive kmers for the plasmids were weakly correlated and covered a wide range of orders, some of which were identified by BAPS-based analysis (Figure 6 for pKJK5 and Figure S5 for pB10 and RP4 plasmids). These kmers were found to be predominantly uncorrelated as only < 0.01% of the kmers had a significant correlation for their pairwise presence/absence pattern (Pearson test), showing the independence of the signals. For pKJK5, kmers linked with the Gram-negative orders of Enterobacterales, Betaproteobacteriales, Xanthomonadales and Sphingomonadales most strongly predicted permissiveness. For RP4 and pB10, besides Pseudomonadales and Cellvibrionales, which were identified by BAPS analysis, Aeromonadales ASVs appeared to contain the predictive kmers (Figure S4), which is in line with recent evidence of HGT for the *Aeromonas* genus in aquatic environments [44]. Predictive kmers, whose absence was linked with permissiveness, had a distinctive distribution for plasmids: For RP4, the kmers for absence predominantly occurred in Flavobacteria strains, which consistently showed a lower permissiveness for the plasmid (Figure S5). For pKJK5, the signals occurred, however, throughout various clades. The kmer analysis appeared to identify further predictive signals.

**Figure 6.**
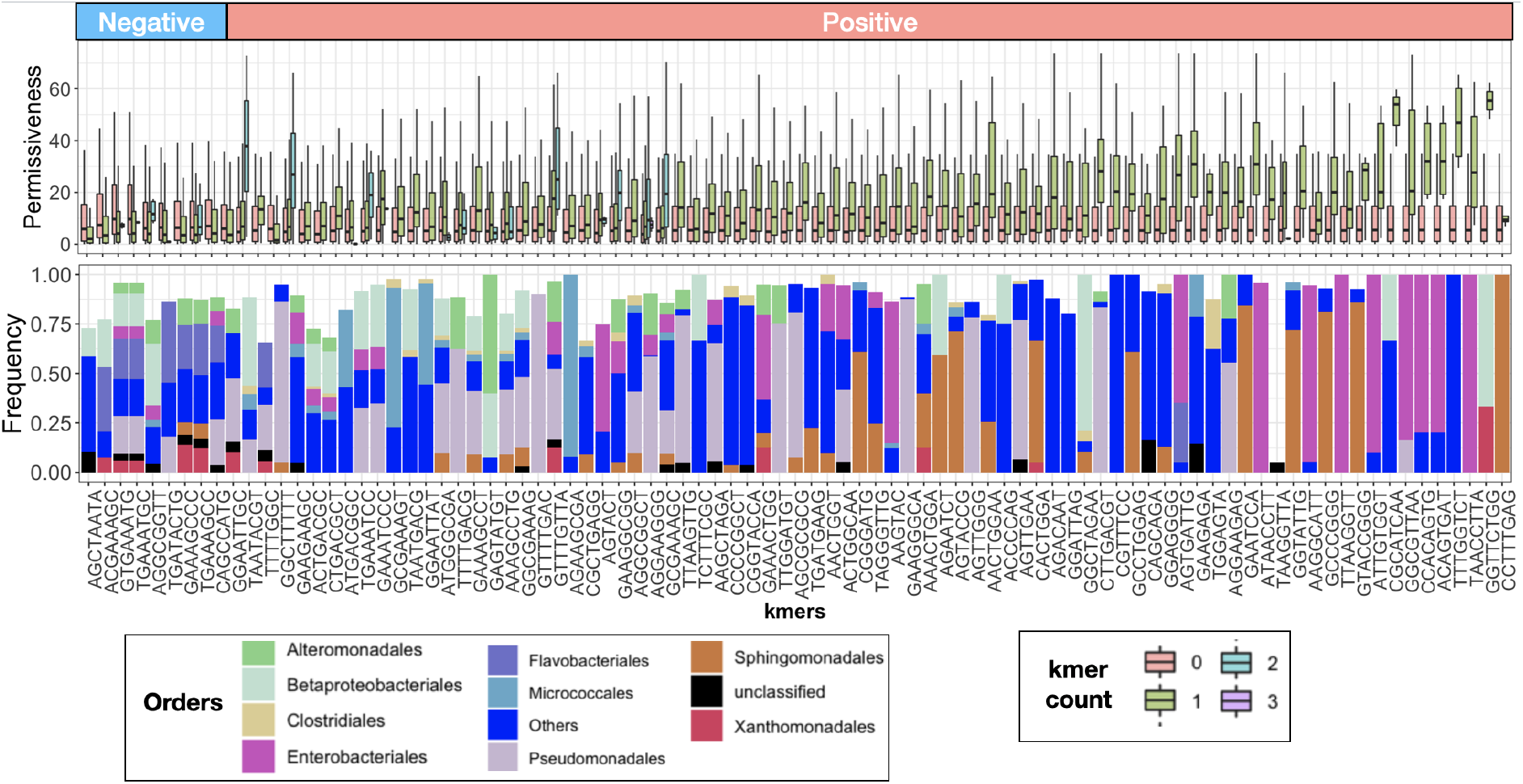
Feature importance analysis for kmer-based predictions. The list of predictive kmers for pKJK5 and the relative frequency of taxa with the respective kmer, showing the effect of the presence of the different counts of the concerned kmers on the plasmid permissiveness. The box plots show the distribution of permissiveness values for ASVs (taxa) bearing the kmer. The groups of box plots are sorted according to the difference in the mean of permissiveness for ASVs with and without the kmer. The negative and positive groups of boxplots are for kmers, the absence and presence of which are linked with increased permissiveness, respectively. The bar plots show the distribution of order memberships for the kmers, showing orders that were enriched in the group of ASVs harboring the kmers compared to the baseline population.

**Figure 7.**
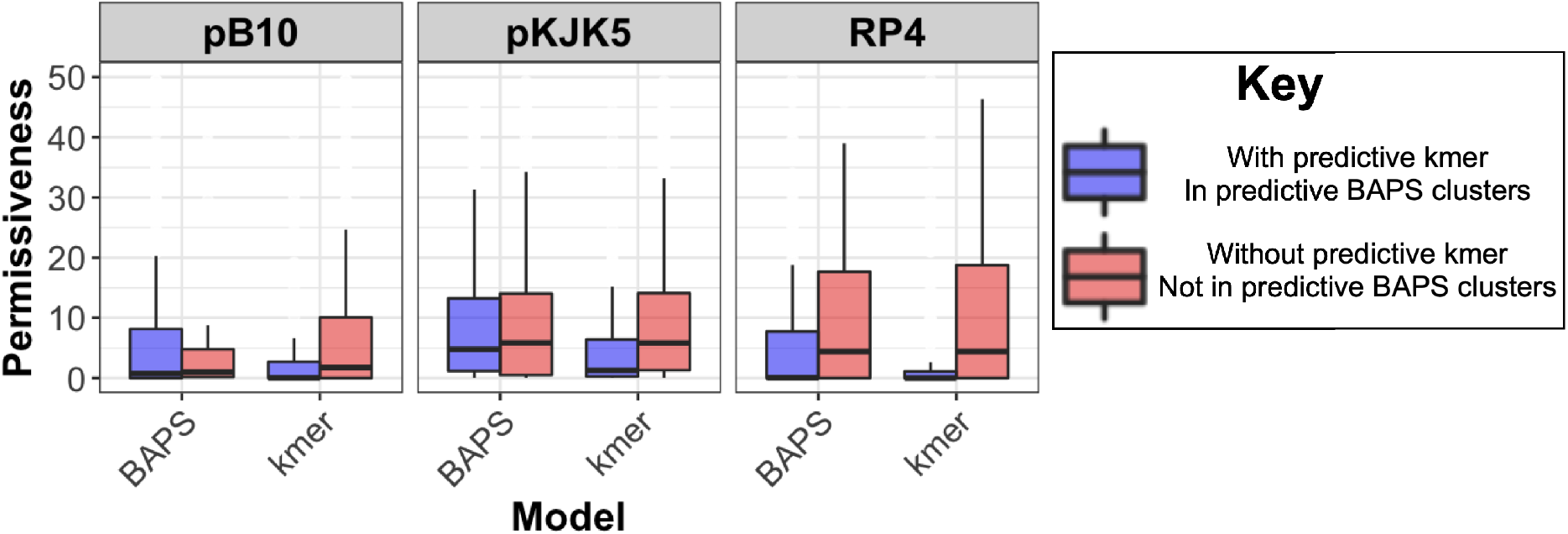
The comparison between the permissiveness values for ASVs within and outside the predictive BAPS clusters and with and without the predictive kmers for the three plasmids.

The lower performance of the models based on taxonomic clusters, as compared with kmer predictions, also suggested that elevated permissiveness may have lineage independent signals, which occur across lineages. To identify these signals, we screened the kmers for association with permissiveness after accounting for the lineage association. In total, we filtered 8, 6 and 5 significant kmers with unique distributions for pKJK5, RP4 and pB10 plasmids linked with elevated permissiveness, respectively (Figure S6). Like the BAPS and kmer results, the distribution of significant kmers differed between the plasmids (Figure S6). For pB10 and RP4, the kmers were found to predominantly occur within Gram-negative clades of Pseudomonadales, Aeromonadales and Cellvibrionales and to a lesser extent in Gram-positive Corynebacteriales strains, in which HGT is recognized to contribute to their pathogenicity and AMR [45]. The significant kmers for pKJK5 were found across a wider range of Gram-negative and Gram-positive species (Figure S6). The presence of these host sequence signals in only very distantly related taxa suggests a convergent evolution of molecular mechanisms for transfer and maintenance of IncP-1 plasmids in Gram-positive and Gram-negative strains [46], which may be discovered by whole genome data analysis.

We then examined the extent to which the number of 16S rRNA sequences used for training and mutation rate would affect the performance of the predictive models. Henceforth, we carried out predictions with simulated 16S rDNA data with different values for the strength of the correlation between the predictive kmer and permissiveness, mutation rate and population size. The results indicate that for a wide range of parameter values for population size and correlation of the predictive kmer with permissiveness, the prediction accuracy for the test data set remained between 0.65-0.75 (Figure S7). With increasing population size, the accuracy of the trained model for the test data did not seem to improve, implying the existence of limits for prediction. The extent of over-fitting, i.e., the difference in accuracy between test and training data, increased with mutation rate (Figure S7). This observation may be explained by the high clonality of populations at low mutation rates, which makes it likely that a taxon from the test dataset falls within the same clade as the training dataset, and thus, making the training data highly predictive of the test data. With increasing mutation rate, although a larger number of predictive signals are available, which improves the accuracy of prediction in the training dataset, the generalizability of the model to the test data declines because of a higher divergence between input sequences in training and test datasets. Despite these, the results indicate that the models attain a high accuracy for a wide range of parameter values.

## Discussion

While it is expected that permissiveness for narrow host-range plasmids should have a clear taxonomic signal that facilitates prediction, this was not clear for broad host-range plasmids. Moreover, there is significant microdiversity [47] between strains with the same full length 16S rRNA sequence and even more so with partial sequences used here, which could make predicting permissiveness rather hopeless. Nevertheless, in this study, we demonstrated that this is possible. We presented a machine learning framework to predict plasmid permissiveness from 16S rDNA amplicon sequencing (V3-V4 hypervariable regions) of recipient and transconjugant pools from filter mating assays inoculated with samples from various compartments of the urban water cycle. Despite the short size of the predictor sequence, our results showed that the host genetic information encoded in 16S rRNA sequences may account for around 50% of the total variance across different resistance plasmids. Furthermore, our analysis identified predictive biomarkers for permissiveness of recipient cells and provided evidence for host preferences and plasmid specificity across multiple lineages.

Although our results with IncP-1 plasmids provided evidence for lineage specificity of elevated HGT rate even for broad host-range plasmids, which depended on plasmid type, we also found a broad phylogenetic distribution of elevated permissiveness, with molecular signals spanning multiple divergent clades. Permissiveness presumably requires suitability of hosts for establishing transfer, avoiding entry exclusion and host immunity systems, e.g., restriction-modification and CRISPR-Cas systems [48, 49], and compatibility with hosts and any resident plasmids [50]. Permissiveness also requires expression of plasmid genes to replicate and partition the plasmid for at least a few divisions.

Several studies reported specific genetic conflicts between chromosomal and plasmid genes that lead to high plasmid fitness costs that could be mitigated by mutations in the host chromosome or the plasmid or both [51–57]. This study identified hubs for conjugation that may be due to certain lineages harboring suitable compensatory mutations, thus providing predictive genomic signals for permissiveness. While the taxonomic distribution of these mutations in full genomes are yet to be elucidated, the elevated permissiveness across distantly related taxa points to possible convergent evolution for plasmid uptake and carriage within hosts.

The unexplained variance in the output of the models likely has various causes. The physiology of the host cell at the time of the experiment will depend not only on its genome (its potential physiology), which is not fully represented by its 16S rRNA phylotype, but also on its transcriptome, proteome and metabolome (its current physiology). The latter will depend on the environmental conditions experienced over the last few generations while the host cell was being transported through several water cycle compartments or resided in one particular compartment with longer solid residence times. The filter mating conditions can also affect cellular activities and shift the community composition, e.g., the synthetic wastewater medium is different from the actual water of the sample’s origin. Therefore, the experimental results may not directly apply to the *in situ* plasmid permissiveness. However, standardized test conditions are essential to eliminate environmental confounders. Certainly, seasonal, diurnal and higher frequency temporal fluctuations combined with considerable spatial heterogeneity will affect the microbial community in the various compartments of the water cycle sampled [58]. To minimize the effect of the environmental heterogeneity and host cell diversity on the performance of the predictive models, a more complete, representative and balanced training collection is essential. An ideal sampling framework would include sufficient samples to represent temporal and geographic variation across all compartments, including the different types of wastewater treatment processes. Such a collection would maximize the viability and generality of the trained models.

Several previous studies have attempted to predict complex bacterial traits from full genomic data of naturally occurring strains. The traits included environmental niches and host phenotypes, host specialism, antimicrobial resistance, and bacterial growth features [24, 59, 60]. However, no machine learning model has been developed for HGT. The predictive power of partial 16S rRNA gene sequences with signatures of lineage dependence that we reported here, sets the stage for future research, where full genome sequences or metagenomic features are used as further predictor signals for the models and feature importance analysis. The models developed here and future models can serve as useful tools for assessing the potential risk of mixing different wastewaters containing resistant and sensitive bacterial strains or releasing these waters into receiving aquatic habitats, enabling subsequent resistance transmission across the environmental systems. Tools with such predictive capabilities can play a major role in supporting One Health efforts. The models, trained on data from studies aiming to reduce plasmid transfer in laboratory settings [61], can enable prediction of the effect of such interventions on a wide range of microbial communities in different environments to assess the utility of these interventions in relevant clinical and non-clinical settings.

## Acknowledgements

This work was supported by funding from the Joint Programming Initiative on Antimicrobial Resistance and the Danish Innovation Foundation (JPI-AMR; DARWIN Project 7044-00004B). D.M. was supported by the Joint Programming Initiative on Antimicrobial Resistance (JPIAMR) as part of the TransPred project via MRC grant MR/R004501/1. M.B. was supported by a UKRI Future Leaders Fellowship [MR/V027204/1].

## Supplementary Figures

**Supplementary Figure S1.**
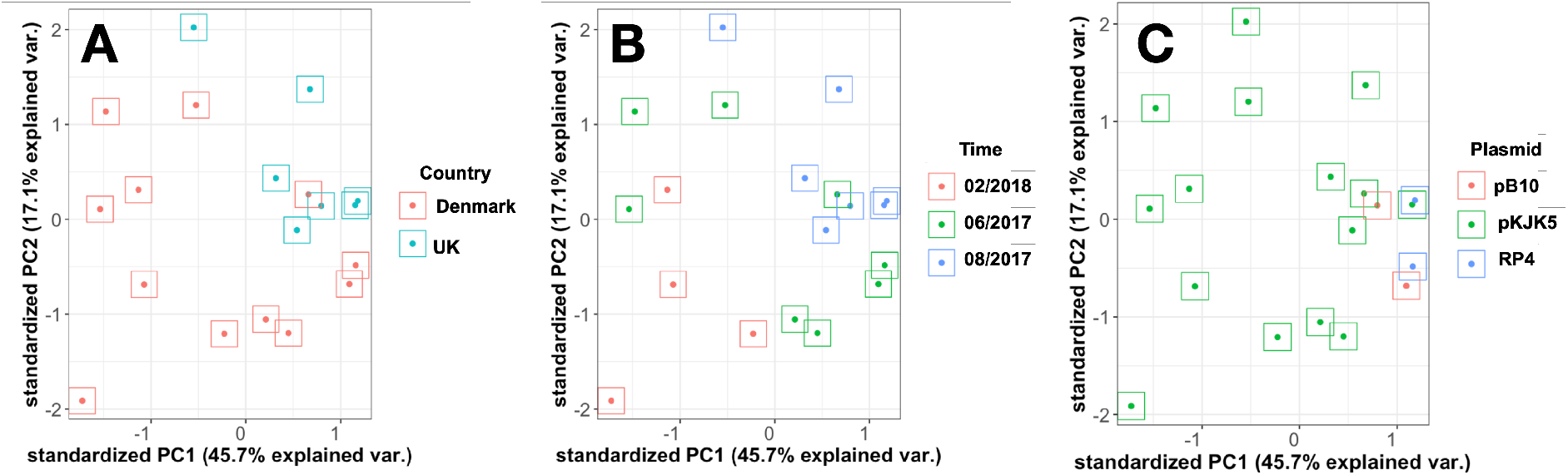
PCA plot for the measured permissiveness values colored according to **(A)** ‘country’, **(B)** time of sampling (month and year) and **(C)** plasmid.

**Supplementary Figure S2.**
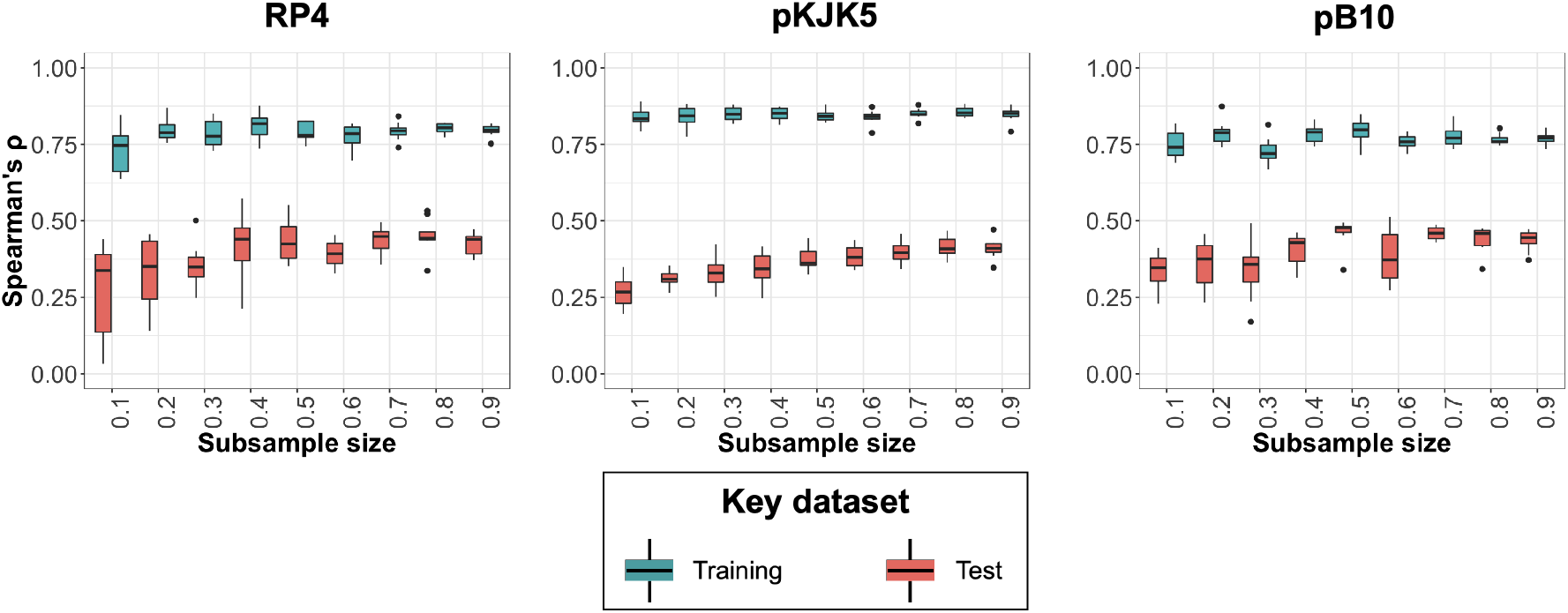
The effect of sample size (transconjugant numbers) on the accuracy of prediction and the extent of over-fitting. Bluegreen and red boxes correspond to the accuracy of prediction for 10 randomly drawn sub-samples for the test and training datasets, respectively. We trained and tuned a random forest model.

**Supplementary Figure S3.**
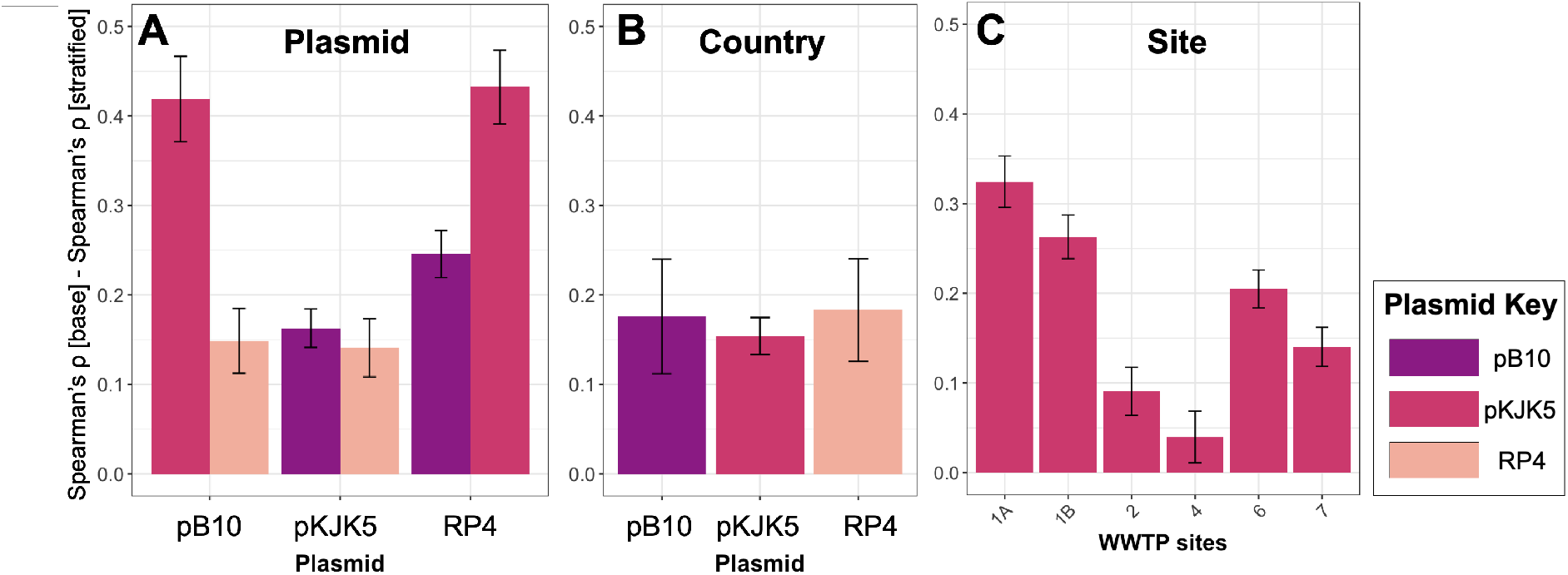
Differences between the accuracy of the baseline model and models trained or tested on stratified training datasets. Higher bars indicate a more significant effect of training sample composition (stratified by site, country of isolation, and plasmid) on prediction accuracy. **(A)** The bars show the difference between the accuracy for the baseline model and models trained on data for one plasmid, as indicated on the x axis, that were then used to predict the permissiveness for the two other plasmids, as indicated by the color key. Here the baseline is the model trained and tested on the dataset for the same plasmid. **(B)** The bars show the difference between the baseline model and models trained on UK data, which were tested on data from Denmark. The baseline model is the model trained on data from both UK and Denmark and tested on data from Denmark. **(C)** The bars show the difference between the baseline model and models trained on pKJK5 permissiveness values for transconjugants from all sites, except the site indicated on the axis that was used for testing. The baseline model is the model trained on data that consisted of transconjugants from all sites. Numbers on the x-axis denote the sites in Figure S1A. The error bars show 95% confidence intervals for ten random training/test data splits.

**Supplementary Figure S4.**
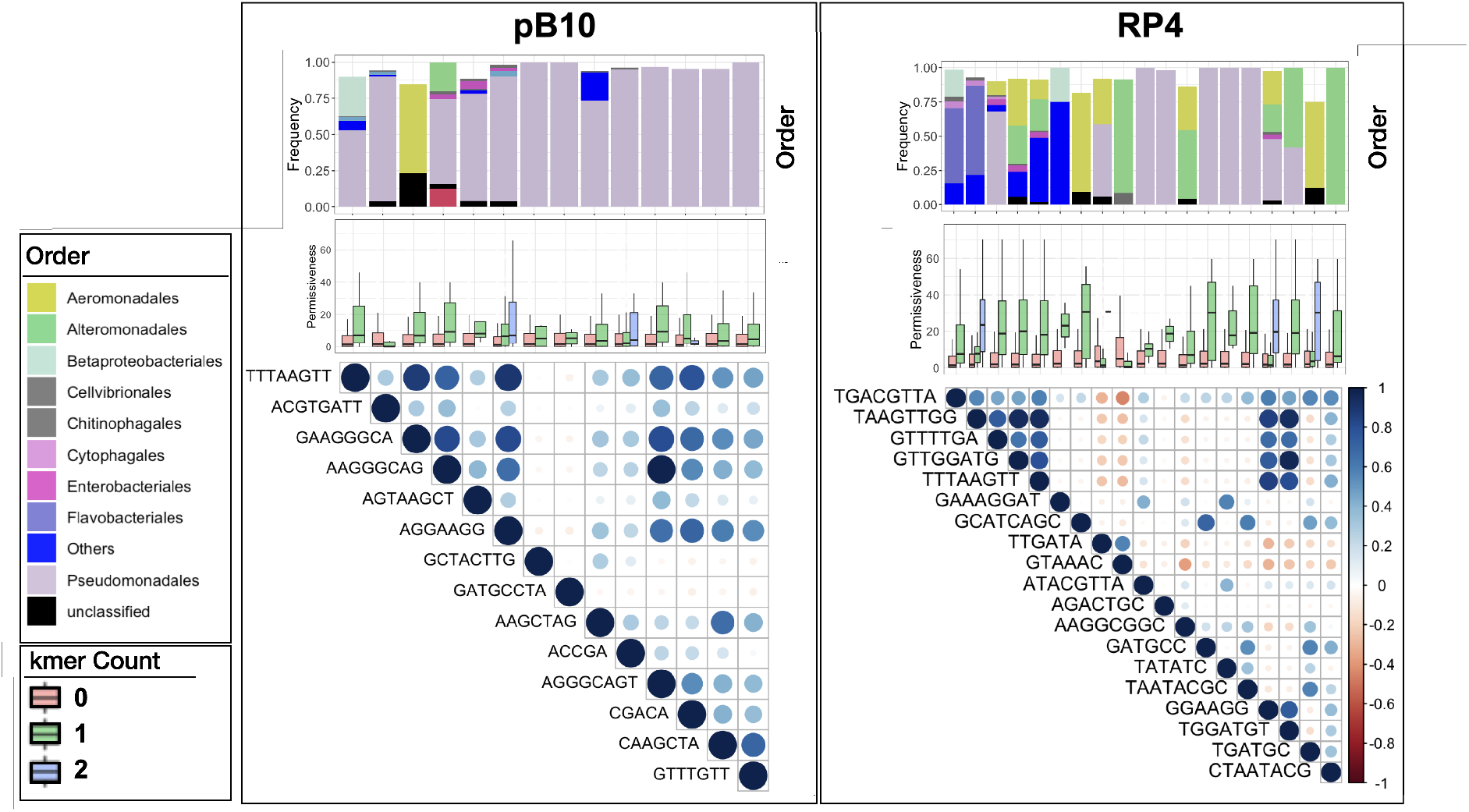
Predictive kmers for **(A)** RP4 and **(B)** pB10 plasmid permissiveness. The correlogram shows the correlation between the presence/absence pattern of the kmers across taxa. The boxplots show the distribution of permissiveness for the plasmids in taxa with different numbers of kmers. The bar plots show the relative frequency of orders containing predictive kmers. Only orders that were enriched for the kmer are shown.

**Supplementary Figure S5.**
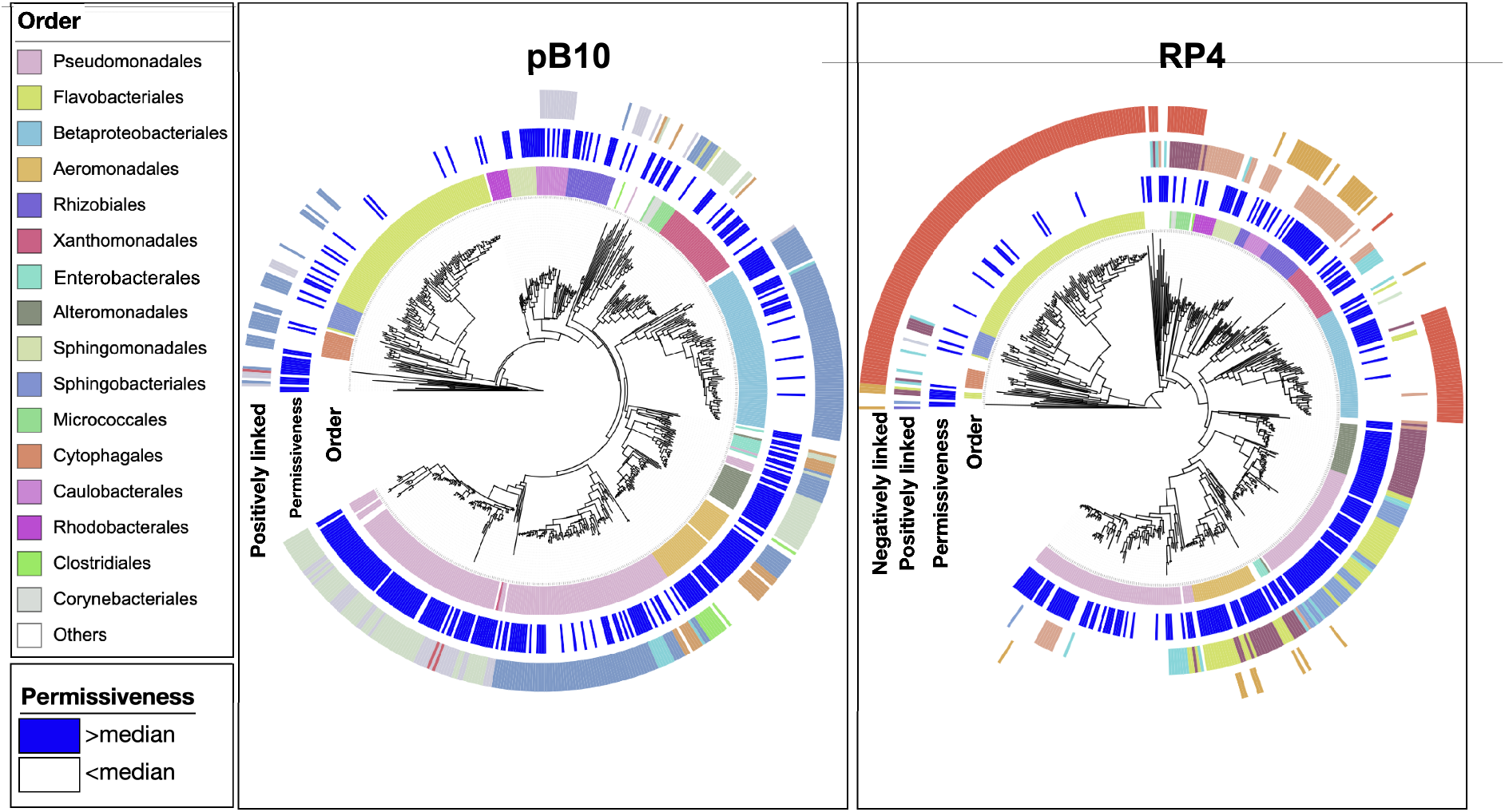
The taxa distribution of kmers in Figure S4, which were positively or negatively linked with permissiveness across the tree. The trees were constructed from distances in the kmer profile for taxa’s 16S rDNA data.

**Supplementary Figure S6.**
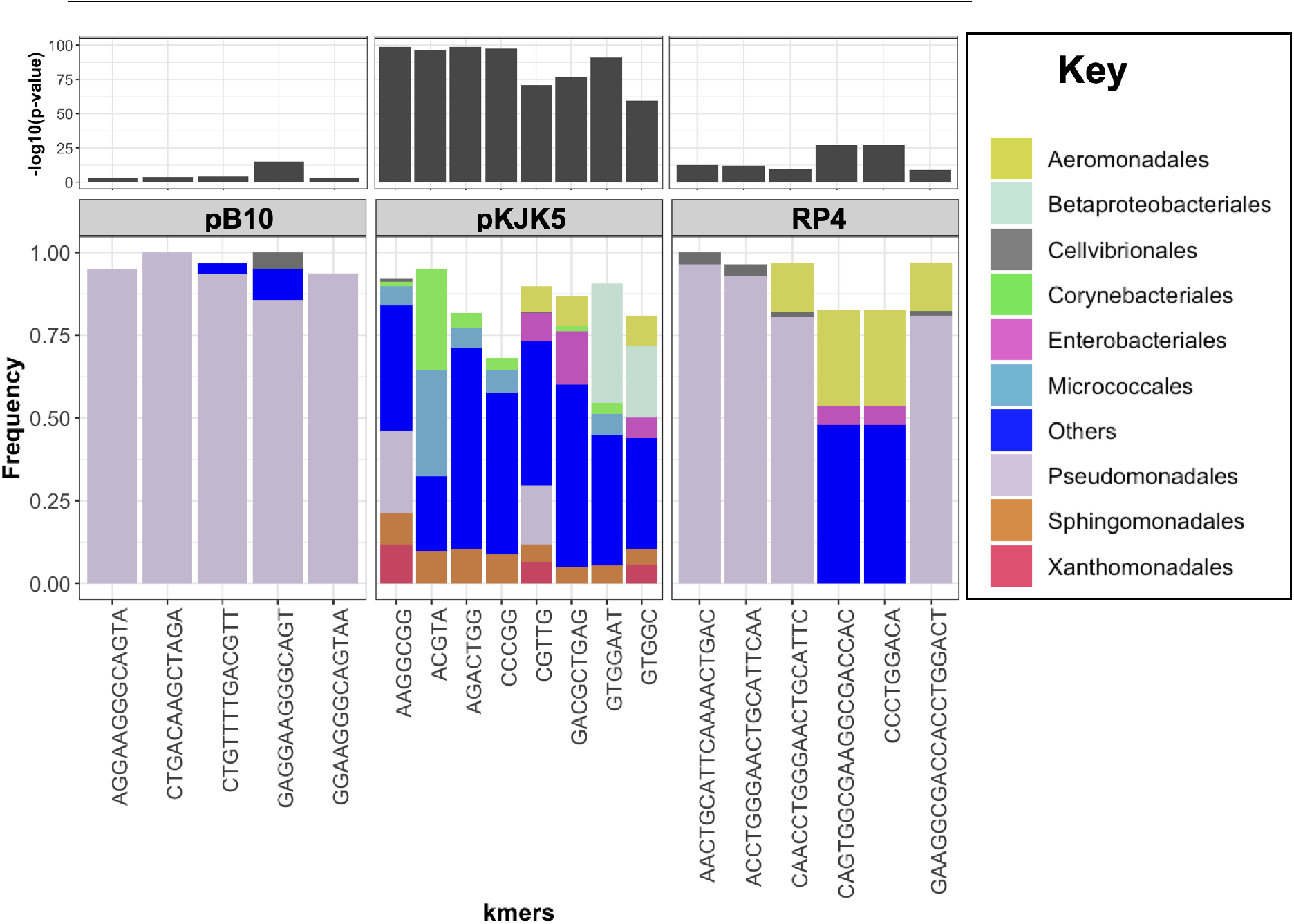
Association analysis for significant kmers that were found after accounting for lineage and clade structure association. Only orders that were overrepresented in the data, i.e., had a higher frequency compared to the baseline frequency in the entire dataset, are shown. The p-value corresponds to the p-value from the association analysis of Scoary, in which the population structure is accounted for.

**Supplementary Figure S7.**
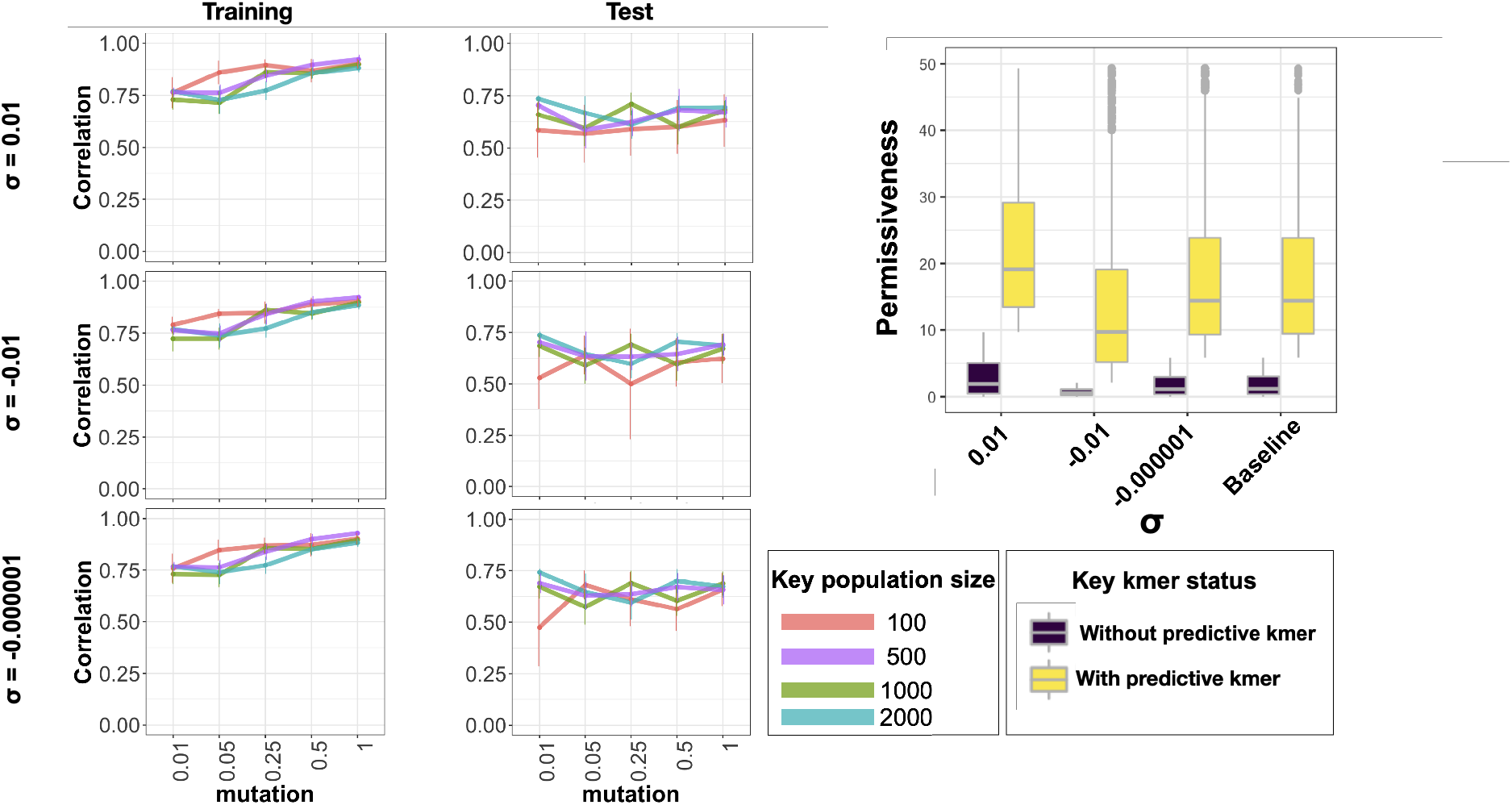
The effect of the number of 16s rRNA sequences used for training, mutation rates and strength of selection on the prediction accuracy, using simulated datasets. Each datapoint corresponds to the average of five simulated populations with the specified values for sigma (strength of selection), population size and mutation rates. The error bars show 95% confidence intervals. The boxplot shows the permissiveness values for ASVs without and with the predictive kmer with different selection strengths (*σ*), which corresponds to permissiveness for the strains bearing the predictive kmer, respectively. The Baseline data shows the permissiveness distribution for pKJK5.

